# Side-chain-directed model and map validation for 3D Electron Cryomicroscopy

**DOI:** 10.1101/014738

**Authors:** Benjamin A Barad, Nathaniel Echols, Ray Y-R Wang, Yifan Cheng, Frank DiMaio, Paul D Adams, James S Fraser

## Abstract

Advances in electron cryomicroscopy allow for the building of *de novo* atomic models into high resolution Coulomb potential maps. While established validation metrics independently assess map quality and model geometry, methods to assess the precise fitting of an atomic model into the map and to validate the interpretation of high resolution features are less well developed. Here, we present EMRinger, which tests model-to-map agreement using side-chain dihedral-directed map density measurements. These measurements reveal local map density peaks and show that peaks located at rotameric angles are a sensitive marker of whether the backbone is correctly positioned. The EMRinger Score can be improved by model refinement, suggesting its utility as an effective model-to-map validation metric. Additionally, EMRinger sampling identifies how radiation damage alters scattering from negatively charged amino acids during data collection. EMRinger will be useful in assessing how advances in cryo-EM increase the ability to resolve and model high-resolution features.

## Introduction

Recent computational and experimental developments in single particle electron cryomicroscopy (cryo-EM) now make it possible, in some cases, to build atomic models without any reference structures^1^. In particular, advances in direct electron detectors^2^, algorithms to classify heterogeneous samples^3,4^, and motion correction^5,6^ are positioning cryo-EM to become a dominant method for determining the structure of dynamic molecular machines^7,8^ and membrane proteins^9,10^. Because these structures are otherwise inaccessible to X-ray crystallography or NMR^11^, it is important to determine the reliability of the resulting atomic models, in particular side chain placement, for their eventual use in directing detailed mechanistic studies or drug development^12^.

All-atom *de novo* cryo-EM models present several unique challenges for validation^13^. First, the Coulomb potential map itself must be validated to ensure that the images are properly recombined and that the resolution estimate is accurate^14^. These validation challenges are primarily addressed by assessing the “gold standard” Fourier Shell Correlation (FSC) between two independently refined half maps^15^. Next the chemical reasonableness of the model is assessed using tools that are commonly applied in X-ray crystallography^16^. Similarly to crystallography, it is essential to balance the agreement to experimental data with the deviation from ideal geometry, while maintaining acceptable stereochemistry, Ramachandran statistics^17^, side chain rotamers^18^, and clash scores^16^.

The weighting between data and prior structural knowledge is key to the third step of model-to-map validation: determining whether the structure is accurately fitted, but not over-fitted, to the map^19^. Several cross validation schemes have been proposed recently^19–21^ and can help to ensure that the model is not only reasonable, but also well fitted to the map. However, real space correlation coefficient-based metrics are dominated by low-resolution, high-signal features and can be complicated by the map B-factor sharpening approaches used prior to model building and refinement^22^. Additionally, these considerations may complicate high resolution model-to-map validation and render it difficult to assess the reliability of the highest resolution features of EM maps, such as side chain or ligand conformations.

A potential solution for assessing the reliability of high resolution models is to examine statistical signatures of the weaker, high resolution, data. In particular, testing whether cryo-EM maps recapitulate the preferred rotameric distributions of protein side chains is particularly appealing since side chains represent the highest resolution features modeled *de novo* by cryo-EM structures. For example, the position of Cγ is constrained to avoid “eclipsed” steric overlaps, predicting that a small map value peak, contributed by the scattering from Cγ, should occur at rotameric χ1 dihedral (N-Cα-Cβ-Cγ) angles near 60°, 180°, and 300° (-60°)^23^. Previously, we have used Ringer^24,25^ to measure the electron density at all possible positions of the Cγ atom for each unbranched side chain under ideal stereochemistry and fixed backbone assumptions. The primary conformation, which is usually well modeled by the crystallographic structure, is defined by a local peak in the distribution of density vs. dihedral angle. In addition, secondary electron density peaks in this distribution can represent alternative side-chain conformations. Across >400 structures, we observed that these secondary peaks were strongly enriched at rotameric positions, which suggested that the secondary peaks represented unmodeled alternative conformations that are populated enough to rise above the noise levels in the electron density map^25^.

Here, we examine whether significant side chain density can be observed in EM maps by measuring the distribution of map value peaks around the χ1 dihedral angle and testing whether the primary peaks are enriched at rotameric positions. Our method, EMRinger, can be used as a global validation metric as structure refinement proceeds and highlights specific areas where manual intervention can be used to improve the local fit of the model. As an additional application, we use EMRinger to probe electron radiation damage to side chains, demonstrating how increased electron dose alters the scattering behavior of negatively-charged side chains. The EMRinger approach directly reveals the side chain information content of EM maps and is complementary to, but independent of, existing validation procedures that report on the resolution of the map, the physical reasonableness of the model, and the detailed fit of the model to the map.

## Results

### Side-chain χ1 map density sensitively reports on backbone positioning

EMRinger interpolates the normalized value of the cryo-EM map at each potential position of the Cγ position around the χ1 dihedral angle, assuming the currently modeled N, Cα, and Cβ atomic positions (**Fig. 1A**). We next plot the distribution of map values by dihedral angle (**Fig. 1B**), which reveals local information about both the map and correctness of the backbone of the atomic model. The peak in the distribution represents the most likely position of the Cγ atomof the side chain, even when it is not immediately obvious “by eye”. Based on steric constraints^26^and data mining from high resolution X-ray structures ^18,27^, we expected that high quality EM maps with well fit backbone models would be enriched in χ1 peaks near the rotameric angles of 60°, 180°, and 300°.

**Figure 1.**
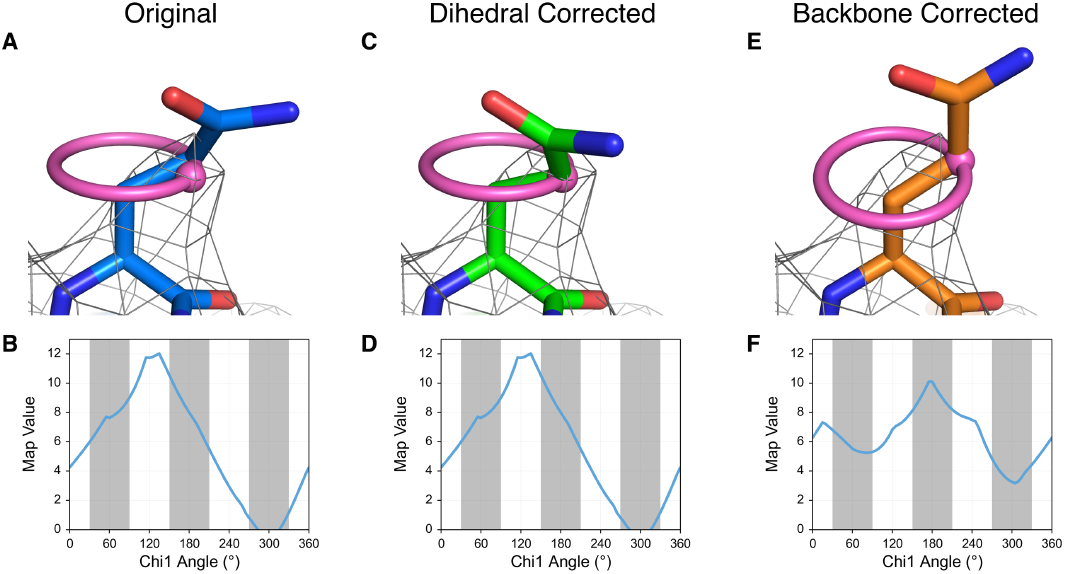
EMRinger χ1 map value sampling reports on backbone position and guides side-chain conformation. (**a**) The side chain of TrpV1 Gln 519 (EMDB 5778, PDB 3J5P) is shown fitted, with a real space correlation coefficient (RSCC) of 0.590, to the potential map, shown at an isolevel of 10. (**b**) The EMRinger scan, reflected by the pink ring in **a**, for Gln 519 of Chain C reveals that the local map value peak (at 130°) occurs at a non-rotameric angle (white bars). This peak, shown as a pink dot in **a**, occurs 30° away from the modeled position. (**c**) The side chain can be rotated so that the χ1 angle is at the map value peak (RSCC = 0.526). (**d**) The EMRinger results are unchanged as the sampling occurs relative to the backbone atoms, which have not moved. (**e**) Alternatively, the backbone position can be corrected with RosettaCM (DiMaio *et al*, Nature Methods, In Press) to place the model near a χ1 map value peak a small reduction on the overall correlation of the residue to the map (RSCC = 0.442). (**f**) The peak at 175° is now in the rotameric region (grey bars).

However, there are several reasons, including noise in the map or an inaccurate model, why a side chain peak might occur at a non-rotameric angle. For example, residue Gln519 of TrpV1^28^(PDB: 3J5P) is modeled in a rotameric position, but has a peak at a non-rotameric angle in a 3.27 Å resolution map (EMDB: 5778) (**Fig. 1A,B**). The distribution in map values by dihedral angle has a single dominant peak, suggesting that there is a local signal above the noise. The lack of a distinct peak can mean that the density threshold is too high, that the backbone is grossly mispositioned, or that the specific area has particularly local low resolution or high noise. However, we observe singular peaks for most side chains in the TrpV1 map, which further suggests that noise is not the dominant reason why the peak occurs in a non-rotameric position. Alternatively, a peak in a non-rotameric position can indicate that the model is incorrect. If the N, Cα, and Cβ atoms are improperly positioned in the strong potential surrounding the backbone, EMRinger will measure the map values in the wrong locations. It is important to note that the side chain is already modeled as rotameric and that changing the modeled side chain dihedral angle does not affect the result of EMRinger because the measurement relies only on the positions of the backbone and Cβ atoms (**Fig. 1C,D**). In contrast, a small backbone adjustment places the Cγ in the map value peak, while maintaining a rotameric side chain model, excellent stereochemistry, and a good map correlation (**Fig. 1E,F**). Thus, EMRinger can identify well-fit backbone models because the local map value peaks will fall at rotameric angles. Our examination of EMRinger plots from several maps suggested that the enrichment of rotameric map value peaks could be used to assess the fit of the backbone model and the overall quality of the EM map.

### EMRinger Score reports on the overall quality of the model and the map

To test the quality of model to map fit, we quantified the enrichment of EMRinger peaks within 30° of rotameric angles as a function of map value threshold. We recorded the position and map value of the peak for each side chain χ1 angle in the 3.2Å resolution 20S proteasome map (EMDB 5623, PDB 3J9I) and observed that the distribution becomes more sharply peaked as the map value cutoff increases (**Fig. 2A, S1A,B**). At lower thresholds, noise flattens the results, with less enrichment for peaks in rotameric regions. Although rotameric regions are sampled more at higher thresholds, fewer residues have local map value peaks above these thresholds, and noise from counting statistics dominates (**Fig. 2B**). To quantify the relationship between sample size and rotameric enrichment, we used the normal approximation to the binomial distribution to generate a model-length independent validation statistic: the EMRinger score (**Fig. 2C, S2**). For the 20S proteasome, the EMRinger score is maximized at the 0.242 normalized map value threshold and the signal is dominated by 1547 rotameric map value peaks, compared to 555 non-rotameric peaks (**Fig. S3**).

**Figure 2.**
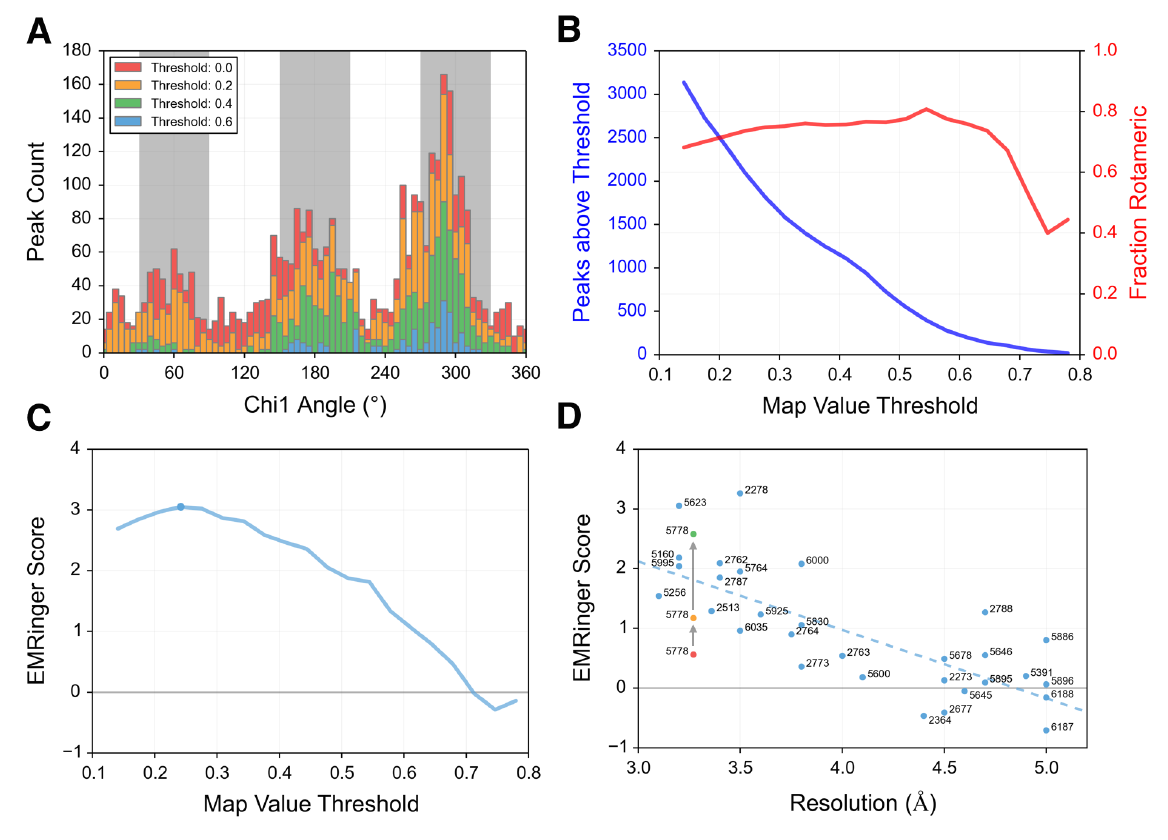
EMRinger reveals statistical enrichment at rotameric χ1 angles in high resolution EM maps. (**a**) Histograms of EMRinger peaks observed above multiple map value thresholds. At high thresholds, more residues are located in the rotameric regions (grey bars). As the threshold lowers, relatively more peaks are added to the non-rotameric regions (white bars). (**b**) Scanning across map value thresholds demonstrates the tradeoff between sampled peaks (left) and fraction of rotameric peaks (right). **(c)** The EMRinger score balances the sample size and the rotameric enrichment and is maximized at a threshold of 0.242 for the proteasome structure (blue circle). (**d**) EMRinger scores for maps deposited in the EMDB with atomic models demonstrate the relationship between model quality and resolution. A linear fit (R^2^ = 0.549) highlights how refinement of TrpV1 improves from the deposited model (red, PDB 3j5p), the transmembrane domain of the deposited model (orange), and a model refined by RosettaCM (green, PDB 3J9J) (DiMaio *et al,* Nature Methods, In Press).

Next, we sampled a series of cryo-EM maps deposited in the EMDB, spanning from 3-5 Å resolution, with atomic models built into the map density (**Fig. 2D**, **Table S1**). The top scoring maps have scores above 3.0: the T20S proteasome, which used a crystallographic model with minimal refinement with MDFF^6^, and the hepatitis B viral capsid, which was built *de novo* and refined using real space refinement in Phenix^29^. Both maps are consistently better than 3.5 Å local resolution^30^, likely reflecting the underlying rigidity of the complexes. Recent mammalian ribosome structures^7,31^, which are dynamic and have more variability in resolution, used masking to reconstruct the highest resolution regions. Refmac reciprocal-space refinement of *de novo* atomic models of these components results in EMRinger scores above 1.85^22^.

The EMRinger approach confirms the resolution dependence of side chain signals, with a strong correlation between decreasing resolutions and decreasing scores (**Fig. 2D**). Since a random distribution should produce an EMRinger score of 0, the trend line suggests that the χ1 angle of side chains can be resolved at 4.5 Å resolution or better. We observed similar trends in decreasing EMRinger score as maps of the T20S proteasome were progressively low-pass filtered (**Fig. S4**). These results demonstrate how the EMRinger score quantifies the standard visual check that side chains are resolved in high-resolution maps, providing insight into the quality of the high resolution features of the map and the model.

### EMRinger score is highly sensitive to improvements during refinement

A notable exception to the trend of increasing score with higher resolution is TrpV1^28^ (**Fig. 2D**), which had a low EMRinger score (0.56) despite high resolution map (3.27 Å). This *de novo* model was built manually and not subjected to either real-or reciprocal-space refinement. Upon exclusion of the poorly resolved ankyrin domain, the EMringer score increases to 1.17, as only the atoms modeled into the highest resolution data remain (**Fig. S1C, Table S1)**. This suggests that atomic models may be more appropriate for the high resolution transmembrane region than for the ankyrin domain. Further rebuilding and refinement using RosettaCM (DiMaio *et al*, Nature Methods, In Press) gradually improved the EMRinger score in most trials (**Fig. 3A**). Multiple refinement trajectories led to consistent improvements in EMRinger score from 1.17 to above 1.75. The best RosettaCM trajectory improves the EMRinger score to 2.58, while the validation metrics for an independent reconstruction improve by a small margin (**Fig. 3B**, **S5**, **Table 1**). In contrast to existing measures, such as real-space correlation or FSC, the EMRinger score is sensitive to features at lower map values, amplifying improvements in the model that only show a minor impact in the agreement-to-density term used by RosettaCM. Consistent with the overlap between the geometrical and conformational components of the Molprobity score and the Rosetta energy function, refinement also improves MolProbity scores dramatically (**Table 1**).

**Figure 3.**
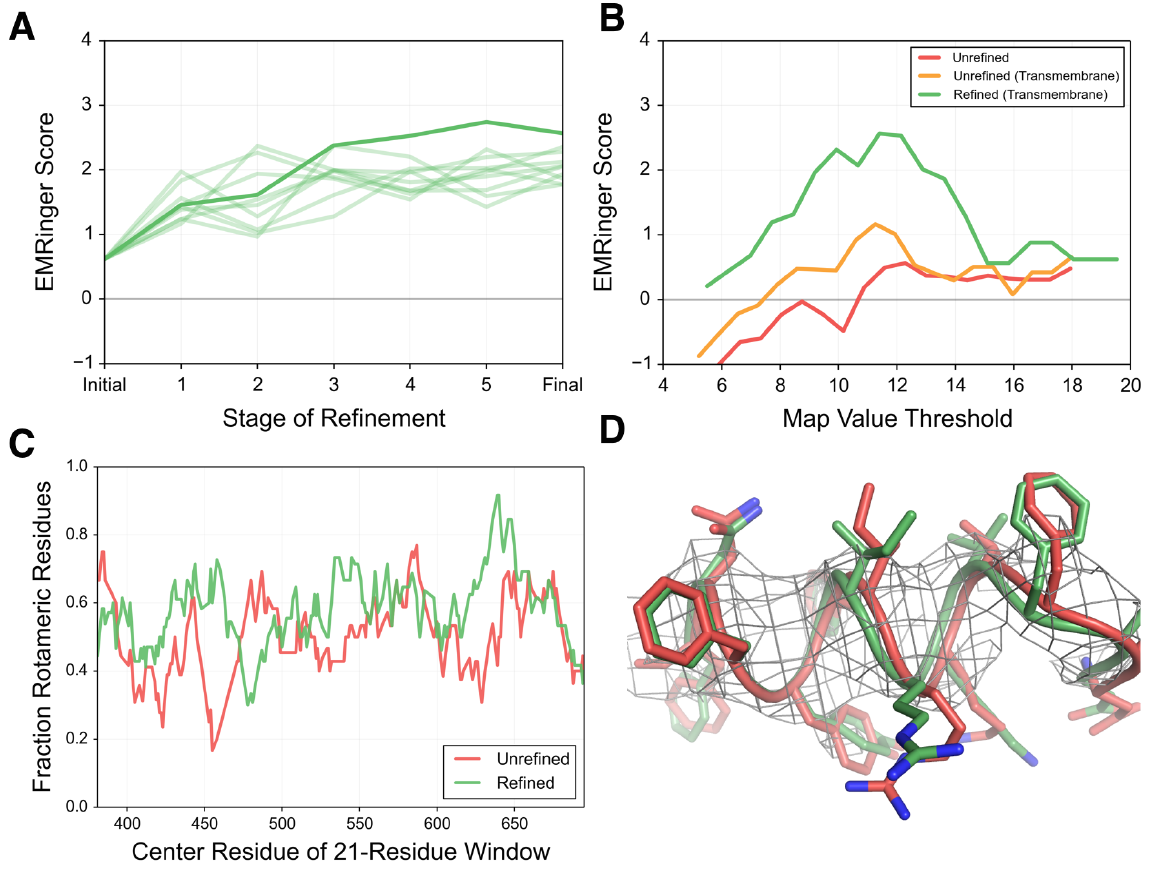
EMRinger Scores report on effective refinement of atomic models into EM maps. (**a**) The EMRinger improves during refinement. RosettaCM (DiMaio *et al,* Nature Methods, In Press) trajectories for 9 trials are shown in light green with the final refinement shown in dark green. (**b**) Map value threshold scan for the unrefined model of TrpV1 (red, EMDB 5778, PDB 3J5P), the transmembrane region of the deposited TrpV1 model (orange), and for the model of TrpV1 refined by RosettaCM (green, PDB 3J9J) show the improvement during refinement. (**c**) Analyzing the unrefined (red) and refined (green) models in the transmembrane region highlights how portions of the model experience dramatic increases in rotameric peaks after refinement. (**d**) The unrefined (red) and refined (green) TrpV1 models are shown in density (isolevel of 10), revealing that small shifts in the placement of backbone of the alpha helix improves EMRinger statistics.

**Table 1.** Statistics pre-and post-refinement. Cross correlation, FSC_mas_k, MolProbity scores and EMRinger score are calculated for the full unrefined TrpV1 model (EMDB 5778, PDB 3J5P), the transmembrane domain of the unrefined model, an intermediate model during refinement of the transmembrane region, and the final refined transmembrane region.

|  | **Unrefined** | **Unrefined**<br>**(Transmembrane**<br>**Region)** | **Refinement Step 2**<br>**(Transmembrane**<br>**Region)** | **Refinement Final**<br>**(Transmembrane**<br>**Region)** |
| --- | --- | --- | --- | --- |
| **CC**<br>**(3.27 Å Cutoff)** | 0.676 | 0.726 | .715 | 0.728 |
| **CC (Training**<br>**Map)** | 0.663 | 0.715 | 0.708 | 0.718 |
| **CC (Testing**<br>**Map)** | 0.664 | 0.714 | 0.705 | 0.713 |
| **Integrated**<br>**Model-Map**<br>**FSC**<br>**(15-3.4 Å)** | 0.473 | 0.553 | 0.513 | 0.526 |
| **All-atom**<br>**Clashscore**<br>**(MolProbity)** | 77.90 | 100.78 | 2.32 | 2.09 |
| **Modelled**<br>**Rotamer**<br>**Outliers**<br>**(MolProbity)** | 26.6% | 30.94% | 0.35% | 0% |
| **EMringer Score** | **0.56** | **1.17** | **1.61** | **2.58** |

To identify the local changes responsible for these improvements, we analyzed 21-residue rolling windows along the length of the protein for the percent of peaks that occurred near rotameric angles (**Fig. 3C**). The specific effects of the RosettaCM refinement can be seen in small backbone shifts, which move the C-beta atoms so that the peak value moves into a rotameric position (**Fig. 3D**). These results demonstrate how small corrections of backbone position along secondary structures, introduced through independently-scored refinement procedures, can lead to improvements in EMRinger score and the accuracy of the resulting model.

### EMRinger Score reveals the residue-specific effects of radiation damage

Radiation damage can severely limit the ability draw biological conclusions from EM data^32^. Because the electron beam also induces motion of the sample, the impact of radiation during data collection has been difficult to assess. Recent motion corrected analyses have indicated that high-resolution information degrades as a function of total electron dose, likely as a result of radiation damage^8^, and that the signal in the 5Å shell degrades rapidly in the second half of data collection^6^. In addition to these global metrics, previous work has hypothesized that differential radiation damage causes negatively charged glutamate and aspartate residues to have weaker density than neutral, but similarly shaped, glutamine and asparagine residues^8,33,34^.

To quantify the effect of radiation damage on the high resolution features of the map and to address whether effects vary by residue type, we used EMRinger for dose-fractionated maps of the T20S proteasome. The overall EMRinger score degrades as a function of dose, with a sharp loss of signal beginning around the 15th frame, corresponding to a total dose of ~18 e^-^/Å^2^ (**Fig. 4A**). Next, we performed EMRinger analysis on different subsets of amino acids. Amino acids with charged side chains generally lost signal more quickly as a function of dose than average, whereas aromatic residues were much more resistant to degradation (**Fig. 4A**). Most notably, negatively charged side-chains appeared to lose signal much faster than positively charged side-chains, with EMRinger score dropping to zero by the map centered on the 8th frame.

The divergent results of EMRinger analysis of negatively charged side chains may be in part explained by the differential radiation damage effects that have been previously hypothesized. However, since a map comprised only of noise (in the extreme of radiation damage) should result in a score of zero, this effect is not sufficient to explain negative EMRinger scores observed in later frames. We examined the specific behavior of the negatively charged residues and observed that the initial map value peaks for some negatively charged residues inverted and became a local minimum in later frames (**Fig. 4B, C**). This behavior is in contrast to the flattening effect, where a peak slowly degrades into noise, seen generally for other residue types (**Fig. 4D, E**). The inversion of the peak may result from the electron scattering factors of negatively charged oxygen atoms, which are positive at high resolution but become negative at low resolution^35^. This radiation damage effect would lead to a negative scattering contribution near the true (rotameric) position in subsequent maps. Because the rotameric peak of the original map can therefore be lowered below the baseline, EMRinger will then identify a new peak at a different local maximum in the damaged map. This new local maximum is more likely to occur at non-rotameric angles because the original rotameric angle is now suppressed by negative scattering contributions in the damaged map. The net effect of the negative scattering behavior could therefore result in an enrichment of peaks at non-rotameric positions and, consequently, a negative EMRinger score after significant radiation damage has accumulated.

**Figure 4.**
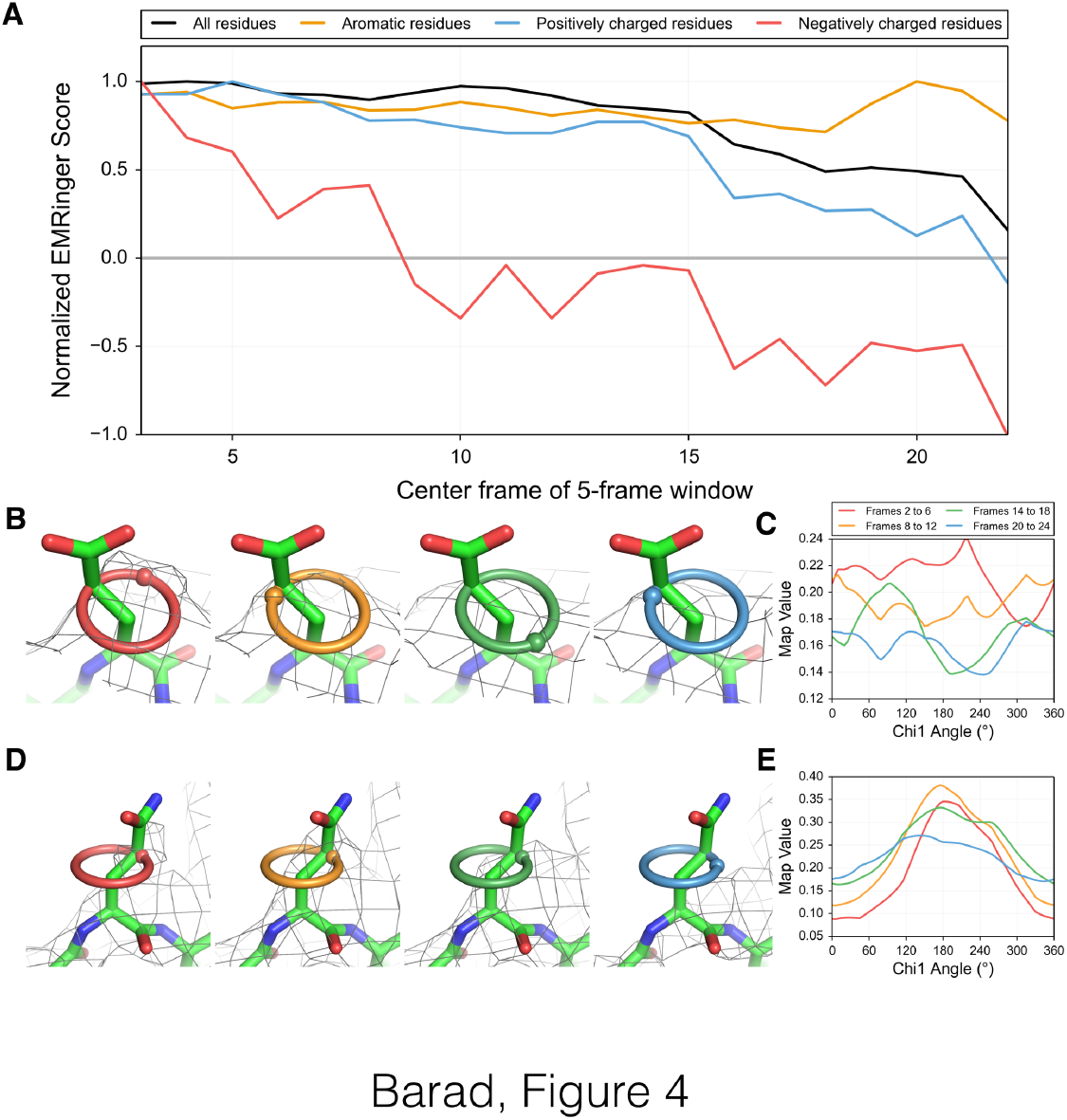
Acidic residues are differentially altered by radiation damage. (**a**) Normalized EMRinger scores are plotted for the T20S proteasome model (PDB: 3J9I) against maps calculated from 5 frames of data. Scores for the entire model (black), the aromatic residues (orange), and the positively charged residues (blue) slowly decrease as a function of dose. In contrast, negatively charged residues (red) experience a rapid drop and fall below a random score of 0. (**b**) Proteasome chain D residue Glu 99 shown in density (isolevel 0.18) for maps generated from frames 2-6 (red ring), 8-12 (orange ring), 14-18 (green ring), and 20-24 (blue ring), with spheres showing local map value peaks. (**c**) EMRinger plots for Glu 99 of Chain D corresponding to the maps in **b** show that peaks immediately flatten and eventually invert after high dose has accumulated. (**d**) Proteasome chain 1 residue Gln 36 shown in density (isolevel 0.3) as in **b**. (**e**) EMRinger plots corresponding to the maps in **d** show a gradual loss of signal as a function of dose.

## Discussion

The dramatic advances in electron cryomicroscopy have created new challenges in building, refining, and validating atomic models. EMRinger extends and complements existing cryo-EM validation procedures at multiple levels. For example, the idea that high resolution features are detectable, confirming the resolution estimate, is quantified by the side chain enrichment. Moreover, the enrichment score tests the fine features of the side chain map density, which intersects with validating the physical correctness of the modeled backbone. While current methods test conformational features independently of agreement with the map, the EMRinger tests these features by querying the model and map together. This procedure is responsive to small backbone corrections that increase the accuracy of the model and the ability to draw mechanistic insights from it.

Our work confirms that side chain detail can be resolved in these maps by quantifying the statistical enrichment of map value peaks at rotameric positions of side chains. Although our analysis was restricted to χ1 angles, similar statistical signatures may extend further out along many side chains. These statistical signatures, which are present in maps determined without model-biased phasing, are a strong indicator that the side chain density that has been identified is predominantly signal rather than noise. Our results confirm that recent advances in data collection, processing, and refinement are increasing the resolvability of atomic features and provide a new metric for assessing the reliability of atomic models generated *de novo* from high resolution cryo-EM maps.

Whereas model-to-map agreement metrics are normally dominated by low resolution features, the EMRinger score reports specifically on statistical signatures in high-resolution data. To validate the model-to-map correctness of atomic models from cryo-EM, refinement should result in EM Ringer scores above 1.0 for well-refined structures with maps in the 3-4 Å range. EMRinger scores can be used in concert with cross validation procedures^21^ and with other measures, such as gold-standard FSC-based resolution^13^ and Molprobity statistics^16^. While it is unlikely that maps with highly variable resolution, generated by imaging more dynamic proteins, will display as much rotameric enrichment as more static molecules, successes in classification of images into different maps representing distinct biochemical states^36^ should be accompanied by increases in EMRinger scores. Similarly, EMRinger scores should quantify improvements in resolvability of atomic features due to improved motion correction algorithms or improved balance between dose and radiation damage during data collection. The results of the EMRinger analysis on dose-fractionated data suggest that reconstructions based on different doses may be required to maximize the resolvability of different sets of side chains, just as different degrees of sharpening are commonly used now during model building.

Additionally, the high sensitivity of EMRinger suggests a natural direction for model-building and refinement. At the resolutions commonly used for model building in EM, there are many closely related backbone conformations that can fit the map density with nearly equal agreement. Given a nearly finalized backbone position, side chains with non-rotameric peaks can be adjusted to fix the Cγ atom in the peak density. Subsequently, the backbone conformation and closure to adjacent residues can be optimized to maintain a rotameric side chain conformation, similar to the inverse rotamer approach used in some protein design applications^37^. This procedure could, in principle, be iterated many times to converge on backbones that are consistent with the map and satisfy the rotameric peak constraints exploited by EMRinger. Similar approaches to quantifying statistical signatures in weakly resolved data may also prove helpful for modeling of non-amino-acid structures at lower resolutions, including glycans and nucleic acids^38,39^.

## Acknowledgements

This work benefited from helpful discussions with David Agard, David Baker, Evan Green, Charles Greenberg, Adam Frost, and Sjors Scheres. B.A.B. is supported by training grant T32GM008284. Y.C. is supported by NIH grants GM082893, GM098672 and GM082250. N.E. and P.D.A. are supported by NIH grant GM063210. J.S.F. is a Searle Scholar, a Pew Scholar, and a Packard Fellow, and is supported by NIH OD009180, GM110580, NSF STC-1231306, and UCSF-SABRE Innovation grants.

## Online Methods

All scripts can be found at https://github.com/fraser-lab/EMRingerandcanberunusingPhenix/cctbx python (version numbers greater than 1894).

### Map Values

We loaded CCP4 formated maps using cctbx and used the map voxel values without further normalization. The wide range of normalization procedures used in constructing these maps explains the large differences in threshold values used for different model-map pairs in our study. However, because EMRinger calculations are based on the relative values of a single map, we can compare EMRinger scores between maps without further normalization.

### EMRinger Map Analysis

EMRinger, as implemented in the Phenix software package^40^, is an extension of the Ringer protocol developed previously^24,25^. We adapted EMRinger to work with real-space maps and to rotate the Cγ atom by increments of 5° around the χ1 dihedral angle (starting at 0° relative to the amide nitrogen). EMRinger calculates and records the map value from a potential map at the position of the atom at each increment using the eight-point interpolation function supplied by Phenix. From this scan, EMRinger records the peak map value and the angle at which it is achieved. EMRinger is available as the *emringer.py* script. Real space correlation coefficients were measured by the *em rscc.py* script.

### EMRinger Score for Validation

We sampled all non-γ-branched, non-proline amino acids with a non-H γ atom, and measured the percent of map value peaks above a given noise-cutoff threshold that are near rotameric (60°, 180°, or 300°) positions. To determine the significance of this distribution, we calculated a Z-Score based on a normal approximation to the binomial distribution. EMRinger repeats this process across a range of map value thresholds, ranging from the minimum peak map value in any scan to the maximum, and returns the highest Z-score calculated in this range. (Equation 1) In order to compare Z-scores between models of different structures, the Z-score is rescaled to the “EMRinger Score” to account for the total number of amino acids in the model (Equation 2).

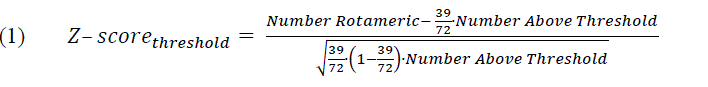

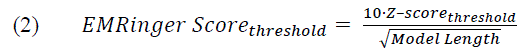

Adjusted EMRinger score does not change when the model and map are multiplied (e.g. in the case of a polymer with high symmetry), so that the score is definitive and no issues arise of how many monomers should be included in the analysis. An EMRinger score of 1.0 sets an initial quality goal for a model refined against a map in the 3.2-3.5Å range, while very high quality models at high resolution generate scores above 2.0. Maps that are highly variable in resolution may have lower EMRinger Scores unless poorly resolved regions of the map are masked out and excluded from the model. Calculation of the EMRinger score is accomplished by the *emringerscore.py* script. Rolling window EMRinger analysis is accomplished by the *emringer_rolling.py* script.

### Refinement of TrpVl with RosettaCM

Refinement of TRPV1 used an iterative local rebuilding procedure to improve local backbone geometry as well as fit to the experimental density data (DiMaio *et al,* Nature Methods, In Press). Refinement began with the deposited PDB structure of TRPV1 (PDB 3J5P). The model was trimmed to the transmembrane region (residues 381-695), and bond angles and bond lengths were given ideal geometry. During local rebuilding, 5 cycles of backbone rebuilding were run; in each cycle, regions with poor fit to density or poor local geometry were automatically identified, and rebuilding focused on these regions. Each rebuilding cycle was followed by side chain rotamer optimization and all-atom refinement with a physically realistic force field. Following this protocol, 1000 independent trajectories were run, and the final model was selected by filtering on two criteria: first, the 80 most nonphysical models were eliminated by assessing each model against the Rosetta all-atom force field; second, fit-to-density was used to rank models and select the best model from these 10.

### Table Statistics

The cross-correlation was calculated using Chimera’s “Fit in Map” tool across all contours and using a resolution cutoff for the calculated map. The integrated FSC was calculated between the model and an independent reconstruction over a masked region covering the protein only. The mask was truncated at 6 Å resolution, and we report the integrated FSC_mask_ over high-resolutions shells only (15 —~3.4 Å). Molprobity statistics were calculated using the validate tool in Phenix nightly build 1894.

### Radiation Damage Analysis

To identify the degradation of map signal with radiation damage, we used EMRinger with a single model across multiple dose-fractionated maps. For each dose-fractionated map, the EMRinger Score is calculated for the model. We calculated additional scores with the amino acids being sampled restricted to different classes (such as acidic or aromatic residues). This residue-specific sampling is accomplished by the *emringer residue.py* script.

## References

1 Liao, M., Cao, E., Julius, D. & Cheng, Y. Single particle electron cryo-microscopy of a mammalian ion channel. Current opinion in structural biology 27, 1–7, doi:10.1016/j.sbi.2014.02.005 (2014).

2 Bai, X. C., McMullan, G. & Scheres, S. H. How cryo-EM is revolutionizing structural biology. Trends in biochemical sciences 40, 49–57, doi:10.1016/j.tibs.2014.10.005 (2015).

3 Lyumkis, D., Brilot, A. F., Theobald, D. L. & Grigorieff, N. Likelihood-based classification of cryo-EM images using FREALIGN. Journal of structural biology 183, 377–388, doi:10.1016/j.jsb.2013.07.005 (2013).

4 Scheres, S. H. Semi-automated selection of cryo-EM particles in RELION-1.3. Journal of structural biology, doi:10.1016/j.jsb.2014.11.010 (2014).

5 Scheres, S. H. Beam-induced motion correction for sub-megadalton cryo-EM particles. eLife 3, e03665, doi:10.7554/eLife.03665 (2014).

6 Li, X. et al. Electron counting and beam-induced motion correction enable near-atomic-resolution single-particle cryo-EM. Nature methods 10, 584–590, doi:10.1038/nmeth.2472 (2013).

7 Brown, A. et al. Structure of the large ribosomal subunit from human mitochondria. Science 346, 718–722, doi:10.1126/science.1258026 (2014).

8 Allegretti, M., Mills, D. J., McMullan, G., Kuhlbrandt, W. & Vonck, J. Atomic model of the F420-reducing [NiFe] hydrogenase by electron cryo-microscopy using a direct electron detector. eLife 3, e01963, doi:10.7554/eLife.01963 (2014).

9 Yan, Z. et al. Structure of the rabbit ryanodine receptor RyR1 at near-atomic resolution. Nature 517, 50–55, doi:10.1038/nature14063 (2015).

10 Lu, P. et al. Three-dimensional structure of human gamma-secretase. Nature 512, 166–170, doi:10.1038/nature13567 (2014).

11 Shi, Y. A glimpse of structural biology through X-ray crystallography. Cell 159, 995–1014, doi:10.1016/j.cell.2014.10.051 (2014).

12 Wong, W. et al. Cryo-EM structure of the Plasmodium falciparum 80S ribosome bound to the anti-protozoan drug emetine. eLife 3, doi:10.7554/eLife.03080 (2014).

13 Henderson, R. et al. Outcome of the first electron microscopy validation task force meeting. Structure 20, 205–214, doi:10.1016/j.str.2011.12.014 (2012).

14 Chen, S. et al. High-resolution noise substitution to measure overfitting and validate resolution in 3D structure determination by single particle electron cryomicroscopy. Ultramicroscopy 135, 24-35, doi:10.1016/j.ultramic.2013.06.004 (2013).

15 Scheres, S. H. & Chen, S. Prevention of overfitting in cryo-EM structure determination. Nature methods 9, 853–854, doi:10.1038/nmeth.2115 (2012).

16 Chen, V. B. et al. MolProbity: all-atom structure validation for macromolecular crystallography. Acta crystallographica. Section D, Biological crystallography 66, 12–21, doi:10.1107/S0907444909042073 (2010).

17 Ramachandran, G. N., Ramakrishnan, C. & Sasisekharan, V. Stereochemistry of polypeptide chain configurations. Journal of molecular biology 7, 95–99 (1963).

18 Lovell, S. C., Word, J. M., Richardson, J. S. & Richardson, D. C. The penultimate rotamer library. Proteins 40, 389–408 (2000).

19 DiMaio, F., Zhang, J., Chiu, W. & Baker, D. Cryo-EM model validation using independent map reconstructions. Protein science: a publication of the Protein Society 22, 865–868, doi:10.1002/pro.2267 (2013).

20 Amunts, A. et al. Structure of the yeast mitochondrial large ribosomal subunit. Science 343, 1485–1489, doi:10.1126/science.1249410 (2014).

21 Falkner, B. & Schroder, G. F. Cross-validation in cryo-EM-based structural modeling. Proceedings of the National Academy of Sciences of the United States of America 110, 8930-8935, doi:10.1073/pnas.1119041110 (2013).

22 Brown, A. et al. Tools for macromolecular model building and refinement into electron cryo-microscopy reconstructions. Acta Crystallographica Section D 71, 136–153, doi:10.1107/S1399004714021683 (2015).

23 Dunbrack, R. L., Jr. Rotamer libraries in the 21st century. Current opinion in structural biology 12, 431–440 (2002).

24 Lang, P. T., Holton, J. M., Fraser, J. S. & Alber, T. Protein structural ensembles are revealed by redefining X-ray electron density noise. Proceedings of the National Academy of Sciences of the United States of America 111, 237–242, doi:10.1073/pnas.1302823110 (2014).

25 Lang, P. T. et al. Automated electron-density sampling reveals widespread conformational polymorphism in proteins. Protein science: a publication of the Protein Society 19, 1420–1431, doi:10.1002/pro.423 (2010).

26 Zhou, A. Q., O’Hern, C. S. & Regan, L. Predicting the side-chain dihedral angle distributions of nonpolar, aromati., and polar amino acids using hard sphere models. Proteins 82, 2574–2584, doi:10.1002/prot.24621 (2014).

27 Shapovalov, M. V. & Dunbrack, R. L., Jr. A smoothed backbone-dependent rotamer library for proteins derived from adaptive kernel density estimates and regressions. Structure 19, 844–858, doi:10.1016/j.str.2011.03.019 (2011).

28 Liao, M., Cao, E., Julius, D. & Cheng, Y. Structure of the TRPV1 ion channel determined by electron cryo-microscopy. Nature 504, 107–112, doi:10.1038/nature12822 (2013).

29 Yu, X., Jin, L., Jih, J., Shih, C. & Zhou, Z. H. 3.5A cryoEM structure of hepatitis B virus core assembled from full-length core protein. PloSone 8, e69729, doi:10.1371/journal.pone.0069729 (2013).

30 Kucukelbir, A., Sigworth, F. J. & Tagare, H. D. Quantifying the local resolution of cryo-EM density maps. Nature methods 11, 63–65, doi:10.1038/nmeth.2727 (2014).

31 Greber, B. J. et al. The complete structure of the large subunit of the mammalian mitochondrial ribosome. Nature 515, 283–286, doi:10.1038/nature13895 (2014).

32 Glaeser, R. M. Limitations to significant information in biological electron microscopy as a result of radiation damage. Journal of ultrastructure research 36, 466–482 (1971).

33 Bartesaghi, A., Matthies, D., Baneijee, S., Merk, A. & Subramaniam, S. Structure of beta-galactosidase at 3.2-A resolution obtained by cryo-electron microscopy. Proceedings of the National Academy of Sciences of the United States of America 111, 11709–11714, doi:10.1073/pnas.1402809111 (2014).

34 Fioravanti, E., Vellieux, F. M., Amara, P., Madern, D. & Weik, M. Specific radiation damage to acidic residues and its relation to their chemical and structural environment. Journal of synchrotron radiation 14, 84–91, doi:10.1107/S0909049506038623 (2007).

35 Mitsuoka, K. et al. The structure of bacteriorhodopsin at 3.0 A resolution based on electron crystallography: implication of the charge distribution. Journal of molecular biology 286, 861-882, doi:10.1006/jmbi.1998.2529 (1999).

36 Fernandez, I. S. et al. Molecular architecture of a eukaryotic translational initiation complex. Science 342, 1240585, doi:10.1126/science.1240585 (2013).

37 Havranek, J. J. & Baker, D. Motif-directed flexible backbone design of functional interactions. Protein science: a publication of the Protein Society 18, 1293–1305, doi:10.1002/pro.142 (2009).

38 Cowtan, K. Automated nucleic acid chain tracing in real time. IUCrJ 1, 387–392, doi:10.1107/S2052252514019290 (2014).

39 Terwilliger, T. C. Rapid model building of alpha-helices in electron-density maps. Acta crystallographica. Section D, Biological crystallography 66, 268–275, doi:10.1107/S0907444910000314 (2010).

40 Adams, P. D. et al. PHENIX: a comprehensive Python-based system for macromolecular structure solution. Acta crystallographica. Section D, Biological crystallography 66, 213–221, doi:10.1107/S0907444909052925 (2010).

